# Hindbrain boundaries as niches of neural progenitor/stem cells regulated by the extracellular matrix proteoglycan chondroitin sulphate

**DOI:** 10.1101/2023.05.02.539049

**Authors:** Carmel Hutchings, Yarden Nuriel, Daniel Lazar, Ayelet Kohl, Elizabeth Muir, Yuval Nevo, Hadar Benyamini, Dalit Sela-Donenfeld

## Abstract

The interplay between neural progenitor/stem cells (NPSC) and their extracellular matrix (ECM), is a crucial regulatory mechanism that determines their behavior. Nonetheless, how the ECM dictates internal processes remains elusive. The hindbrain is valuable to examine this relationship, as cells in the hindbrain boundaries (HB), which arise between any two neighboring rhombomeres, express the NPSC-marker Sox2 while being surrounded with the ECM molecule chondroitin sulphate proteoglycan (CSPG), in chick and mouse embryos. CSPG expression was used to isolate HB/Sox2+ cells for RNA-sequencing, revealing their distinguished molecular properties as typical NPSCs, which express known and newly-identified genes relating to stem cells, cancer, matrisome and cell-cycle. In contrast, the CSPG-/non-HB cells, displayed clear neural-differentiation transcriptome. To address whether CSPG is significant for hindbrain development, its expression was manipulated in vivo and in vitro. CSPG-manipulations shifted the stem versus differentiation state of HB cells, evident by their behavior and altered gene expression. These results provide novel understanding on the uniqueness of hindbrain boundaries as repetitive pools of NPSCs in-between the rapidly-growing rhombomeres, which rely on their microenvironment to maintain undifferentiated during development.

****SUMMARY:**:** Transcriptomic analysis of hindbrain boundaries revels them to harbor cells with neural progenitor\stem cell properties that rely on local extracellular matrix to maintain their undifferentiated state.

## INTRODUCTION

A common theme during development is morphological organization into compartments comprising of segregated cell populations (Morata and Lawrence, 1978). Those segments constitute the building blocks of the body plan, which dictate regional organization within the tissue. Segregation of cells into defined domains rely on their molecular properties that serve for recognition as well as on the presence of boundaries, which assure cell separation as they organize, proliferate and differentiate (Batlle and Wilkinson, 2012; Dahmann et al., 2011; Irvine and Rauskolb, 2001; Kiecker and Lumsden, 2012). These boundaries were suggested to act as physical barriers to prevent neighboring cells from intermingling, as well as organizing centers that determine the fate of flanking cells by secretion of signals (Kiecker and Lumsden, 2005; Krumlauf and Wilkinson, 2021; Rubenstein et al., 1994). In the developing CNS, two well-defined boundary regions appear to function as organizing centers-the Zona Limitans Intrathalamica (ZLI) and the midbrain-hindbrain boundary (MHB), which orchestrate neuronal fates in the thalamus and midbrain/hindbrain through the secretion of Wnts, FGFs and/or SHH (Guinazu et al., 2007; Lim and Golden, 2007; Rhinn and Brand, 2001).

The embryonic hindbrain is an intriguing model to study compartment boundaries (Moens and Prince, 2002). It undergoes a segmentation process along its rostro-caudal axis, resulting in transitory formation of 7 to 8 repetitive compartments, termed rhombomeres (Rhs). Each Rh has unique gene expression patterns promoting regional-specific fates, differentiation of neurons and production of distinct neural-crest streams (Frank and Sela-Donenfeld, 2019; Parker and Krumlauf, 2020). In between the Rhs, sharp domains arise, which are termed hindbrain boundaries (HBs). Remarkably, the HBs have been found to share specific genes and cellular characteristics; they have a fan-shape morphology, enriched extracellular matrix (ECM) and reduced cell proliferation rate, as opposed to the neighboring Rhs (Heyman et al., 1995; Lumsden and Keynes, 1989). The fact that, unlike Rhs, the repetitive HBs share similar properties, together with their ability to regenerate once removed and their conserved formation in vertebrates, stresses their likely significance (Guthrie and Lumsden, 1991; Pujades, 2020; Wilkinson, 2021). Similar to other compartment boundaries, the HBs have been suggested to act as local organizing centers for the adjacent Rhs, achieved through secretion of various signaling molecules, such as FGF in chick or Wnt and Semaphorins in zebrafish (Mahmood et al., 1995; Pujades, 2020; Riley et al., 2004; Sela-Donenfeld et al., 2009; Terriente et al., 2012; Weisinger et al., 2012).

We have previously found that Sox2, a neural progenitor/stem cells (NPSC) master gene known to regulate the self-renewal and multipotency of NPSCs (Lai et al., 2012; Sarkar and Hochedlinger, 2013), is enriched at HBs of st.16-18HH chick embryo (Peretz et al., 2016). We also found that these Sox2^+^ cells (termed here HB^Sox2^ cells) co-express other NPSC markers and constitute two subgroups; one of slow-dividing cells, which populate the boundary core, and the other of amplifying cells localized near the HB-Rh interface. Furthermore, HB^Sox2^ cells were found to undergo differentiation along the ventricular-mantle axis, but also to provide amplifying Sox2^+^ cells horizontally to Rhs, via cell division. Finally, manipulation of Sox2 led to disorganized neurogenesis in the chick hindbrain. Based on these results we have proposed that HBs of avians act as pools of Sox2^+^ NPSCs to provide proliferating progenitors to the intensely-differentiating Rhs. These findings were later reinforced in the zebrafish hindbrain, where HBs were suggested to act as regions of self-renewing progenitors that later on provide differentiating neurons to the hindbrain (Hevia et al., 2022; Voltes et al., 2019). However, as opposed to the chick embryo, Sox2 expression is not enriched in zebrafish HBs. Nevertheless, the corresponding evidences from chick and zebrafish hindbrains underlines a plausible new role for these unique domains. Yet, the mechanisms governing the preservation of HB^Sox2^ cells as NPSCs in-between the Rhs, are not known.

NPSCs in the sub-ventricular zone (SVZ) of the lateral forebrain ventricles, or in the hippocampal dentate gyrus (DG) (Gates et al., 1995; Mira and Morante, 2020), are found in niches typically constructed with a network of ECM (Faissner and Reinhard, 2015; Kazanis and ffrench-Constant, 2011; Mercier, 2016; Su et al., 2019). A dedicated cross-talk has been suggested to exist between NPSCs and their milieu to regulate the cell’s behavior (Morrison and Spradling, 2008; Walma and Yamada, 2020), which is only partially understood. The hindbrain is a valuable system to illuminate this fundamental cross-talk, as we present here that the HBs of chick and mouse embryos display an intense accumulation of the ECM molecule chondroitin sulphate proteoglycan (CSPG). CSPG, which consists of several secreted or membrane-bound subtypes, is a major ECM component in the CNS, known to play critical roles in development and pathology (Carulli et al., 2005; Dyck and Karimi-Abdolrezaee, 2015). Here we present that CSPG primarily surrounds the Sox2^+^ NPSCs in the HBs and prevents the cells from undergoing pre-mature differentiation in both species. CSPG was also used to separate the HB^Sox2^ cells for RNA-seq analysis, which strongly highlighted the properties of the HB^Sox2^ cells as typical NPSCs, in contrast to the non-HB cells, which demonstrated a marked neural differentiation transcriptome profile. Altogether, we present a novel understanding on the molecular uniqueness of the HB cells as new domains of NPSCs in the developing CNS and demonstrate their dependence on their ECM to stabilize their stem/progenitor cell state.

## RESULTS

### Conserved expression of Sox2 and CSPG at hindbrain boundaries of chick and mouse embryos

HBs of st.16-18 chick embryos are comprised of Sox2+ cells while being enriched with proteoglycans, in particular CSPG (Heyman et al., 1995; Peretz et al., 2016; Weisinger et al., 2012). To determine whether these characteristics are avian-specific or also preserved in mammals, we reviewed hindbrains of chick and mice embryos at equivalent stages (st.18 HH and E10.5, respectively) (Fig. 1A,H). In both species, the expression of Sox2 was significantly higher in HB cells’ nuclei compared to Rhs (Fig. 1B,D,I,K). CSPG was co-expressed at these sites, seemingly surrounding the Sox2^+^ cells, hence indicating that HB cells express a membrane-bound form of CSPG (Fig. 1C,E,F,J,L,M, Fig. S1). Quantification of fluorescent area confirmed that both proteins are enriched at HBs of mouse and chick embryos (Fig. 1G,N). To fully validate the precise co-localization of Sox2 and CSPG, we executed a proximity analysis using IMARIS 3D software, by employing a module built to distinguish CSPG molecules that are expressed adjacent to Sox2 cells out of total number of CSPG signal detected (Fig. 1O). On average, nearly 75% of the identified CSPG molecules were found to be expressed in proximity to Sox2+ cells (Fig. 1P, inclusion criteria-distance ≤5 µm; n=7). The spatial co-expression of both proteins was further evaluated, Z-stack images of chick hindbrain along with Z-position scatter plot showed both markers to be mostly expressed in the ventricular and sub-ventricular zones rather than at the mantle layer of HBs (Fig. 1Q,R).

**Figure 1.**
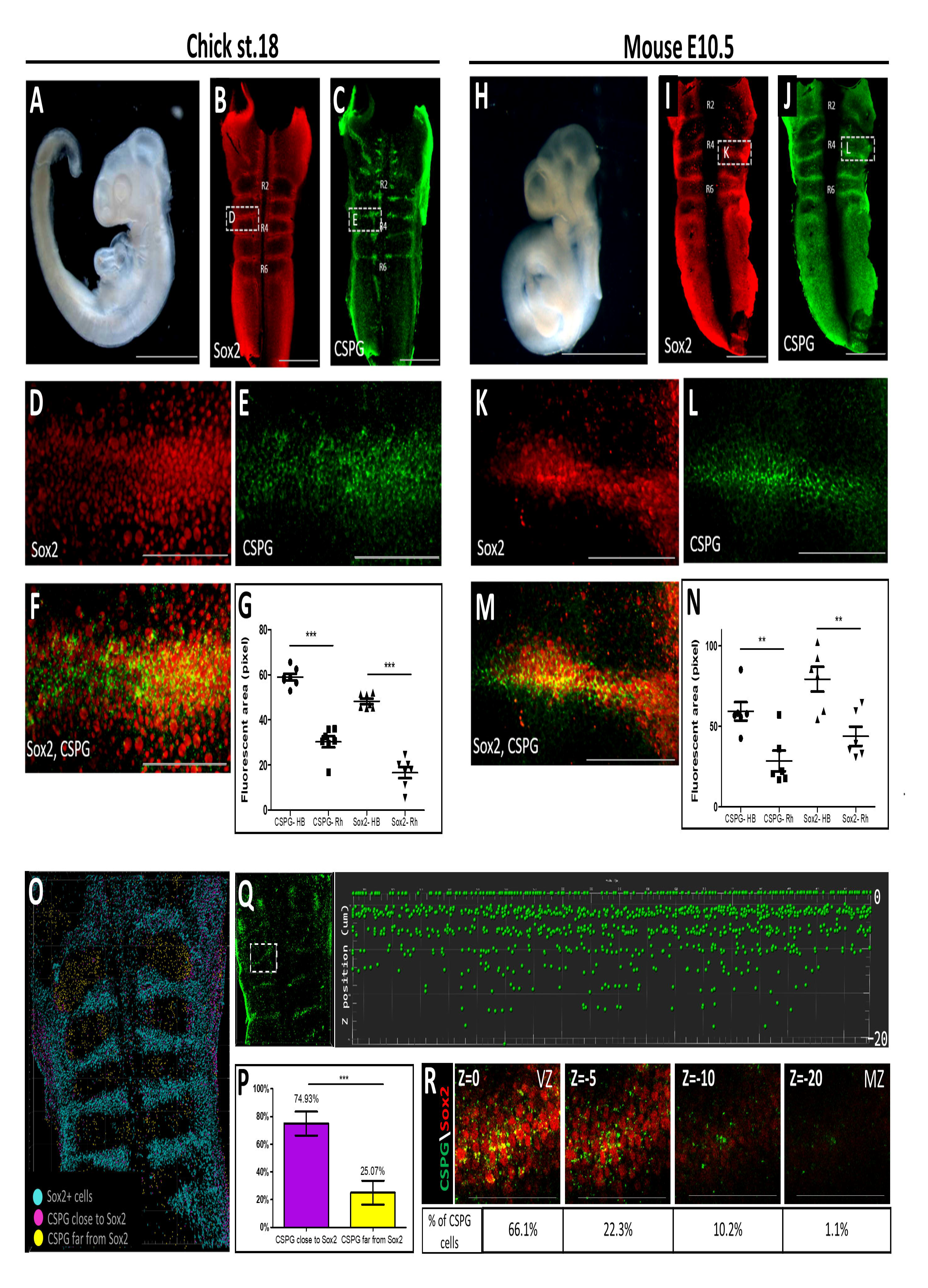
Conserved co-expression of Sox2 and CSPG in chick and mouse hindbrain boundaries. **(A,H):** Representative images of st.18 chick and E10.5 mouse embryos. **(B-F,I-M):** Whole-mounted chick/mouse hindbrains immuno-stained for Sox2 and CSPG. **(G,N):** Quantification of Sox2/CSPG immunostaining at HBs/Rhs in chick or mouse hindbrains. Each dot represents an average of 4 HBs/Rhs in one embryo (n=7). **(O,P):** Representative proximity analysis of Sox2-CSPG signals (n=7, distance ≤ 5 µm). **(Q,R):** Scatter plot (Q) and Z-stack images (R) displaying CSPG distribution at the ventricular-mantle axis of a typical HB region. Boxed area showing region analyzed with Z-distribution presented on the right. Data are mean±s.d. (two-tailed unpaired t-test). ** P< 0.005 *** P<0.0005. Scale bar in A,H= 1000 µm, in B,C,I,J= 100 µm, in D,E,F,K,L,M,R= 50 µm. VZ= ventricular zone, MZ= mantle zone

To further unravel the spatial arrangement of the Sox2^+^/CSPG^+^ HB cells, chick and mouse hindbrains were analyzed by correlative light and scanning electron microscopy on immunofluorescence-stained hindbrains. Intriguingly, in both species the ventricular surface of the hindbrain was uneven, with repeated elevated ridges positioned in-between submerged domains (Fig. 2A,E). This analysis revealed that Sox2 and CSPG accumulate at these elevated zones, verifying them as HBs (Fig. 2B,F). Higher-magnification of these zones revealed their distinct morphology, such as their noticeably larger and smoother apical cell-surface (∼7 µm^2^, Fig. S2; Fig. 2D,H), as opposed to the submerged rhombomeric domains which displayed much smaller (∼1 µm^2^, Fig. S2) and irregular apical surfaces (Fig. 2C,G). These results demonstrate the conserved co-localization of Sox2 and CSPG in the HBs, together with the unique topography of these domains in chick and mouse.

**Figure 2.**
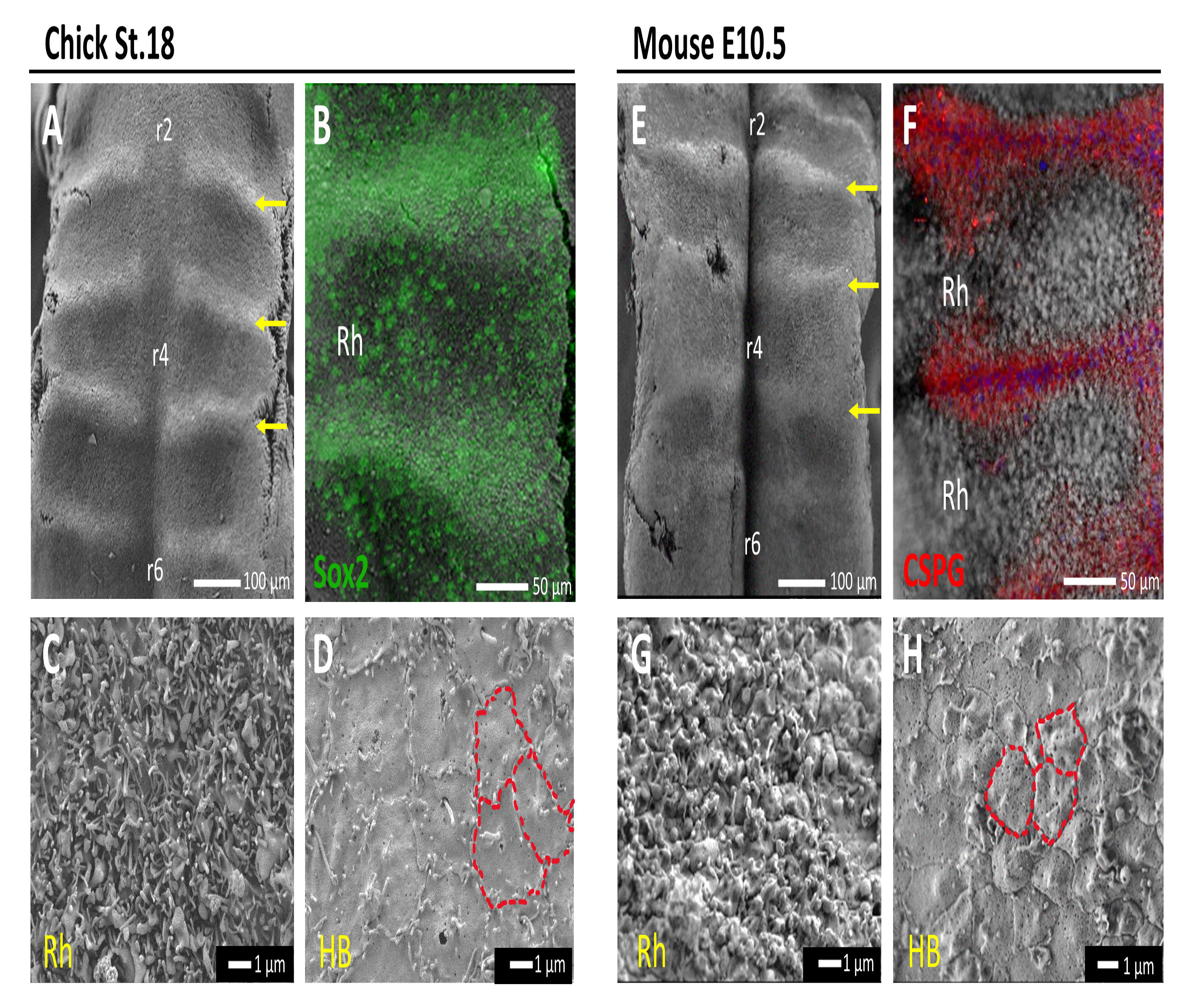
Scanning electron microscopy analysis of chick and mouse hindbrains. **(A,E):** X200 magnification of SEM images of chick and mouse flat-mounted hindbrains, showing HBs as elevated ridges (arrows) and Rhs as grooves. **(C,D):** X10,000 magnification of a representative Rh (C) or HB (D) areas in chick. **(G,H):** X8,000 magnification of a representative Rh (G) or HB (H) in mouse. Red lines mark HB cells. **(B,F):** Correlative light and scanning electron microscopy showing correlation between the elevated HBs regions and Sox2 (B) or CSPG (F) expression.

### Modifications of CSPG levels affects the differentiated state in the hindbrain

To test whether CSPG plays a role in the hindbrain, we knocked-down CSPG by using the procaryote enzyme chondroitinase ABC (ChABC), which digests the CS-chains on CSPGs (Muir et al., 2010). ChABC was either injected into the hindbrain lumen as a soluble protein (mixed into a thermo-sensitive hydrogel (Shriky et al., 2020)), or electroporated as ChABC-encoding plasmid into one side of the hindbrain neuroepithelium (Muir et al., 2010). Initially, the decrease in CSPG levels was confirmed for both treatments by comparing immunostaining in control and treated embryos (Fig. 3A,B). Quantification of the fluorescence levels as well as flow-cytometry analysis of CSPG+ cells, verified the significant reduction in CSPG-expressing cells upon treatment with ChABC (Fig. 3C,D, Fig. S3B), which was not coupled with increased cell death (Fig. S3A). Subsequent analysis of Sox2 expression in the ChABC-treated hindbrains revealed a significant reduction in Sox2 levels compared to controls (Fig. 3A-D). As a reduction in Sox2 expression may shift the cells towards a more differentiated state, we also examined whether expression of the early neural differentiation marker Tuj1 (beta-tubulin III) was altered in the hindbrain. Tuj1 has been previously found to be expressed in the chick hindbrain, mostly accumulating in the mantle zone of HBs with some presence in the Rhs (Peretz et al., 2016). Evidently, treatment with ChABC has led to enhanced expression of Tuj1 along the hindbrain (Fig. 3E,F). Real-time RT PCR analysis further confirmed a reduction in *Sox2* and an increase in *Tuj1* mRNA in the ChABC-treated hindbrains, in agreement with the immunostaining (Fig. 3G). Moreover, flow-cytometry analysis found about 72% increase in Tuj1+ cells in the treated hindbrains (Fig. 3H). The decrease in Sox2 expression and corresponding increase in Tuj1 levels upon loss of CSPG, suggest it may play a role in regulation of differentiation in the hindbrain.

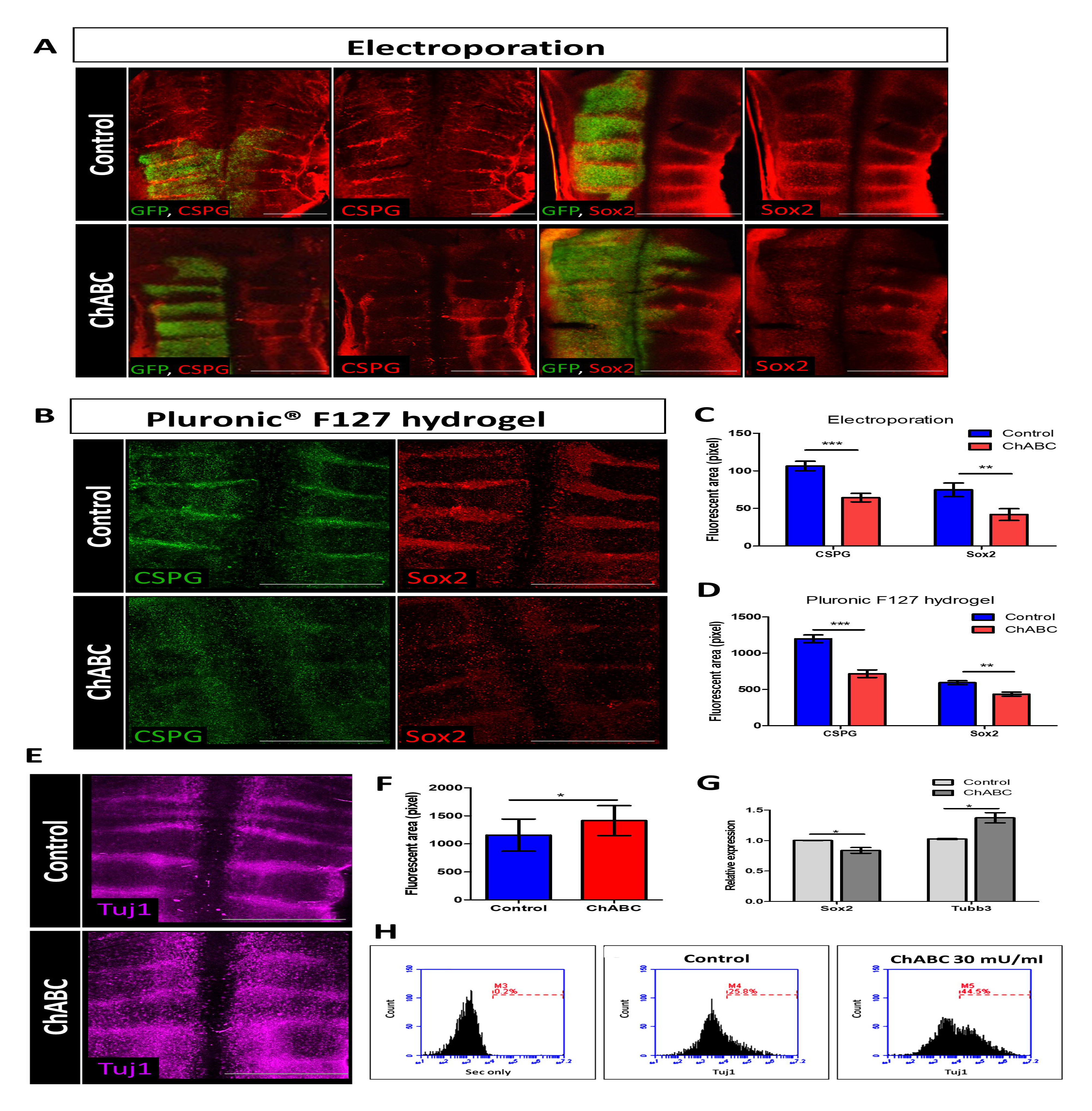

The role of CSPG was next analyzed ex vivo. We have previously demonstrated that hindbrain cells can be grown in primary cultures that display a mixed population of cells; some cells behave as typical cultured NPSCs, as they form floating neurospheres, while others adhere to the surface, create monolayers and extend neurites, as expected from differentiating neurons (Peretz et al., 2018). Hindbrain-derived neurospheres are enriched with Sox2-expressing cells in their core, whereas neural differentiation markers such as Tuj1, MAP2 and 3A10 are evident in the sphere’s periphery. When spheres collapse, they begin to form monolayers with networks of Tuj1/MAP2/3A10-expressing neurites. To evaluate how CSPG removal affects hindbrain-derived cells, primary cultures were prepared from st.18 chick or E10.5 mouse hindbrains and grown in media which support NPSC’s survival (Peretz et al., 2018). Cells were added with ChABC or external CSPG compound, to simulate the effect of loss or excess CSPG (Sirko et al., 2010a). Flow-cytometry analysis revealed a nearly 50% reduction in CSPG+ cells upon ChABC treatment (Fig. S4B), as also detected by immunofluorescence staining (Fig. S4A), confirming its inhibitory activity in vitro.

Next, cells were monitored by live-imaging for 5 days. At the end of the incubation period, each cell group exhibited distinct patterns; Control cells displayed multiple phenotypes including free-floating spheres, adherent spheres and some neurites, ChABC-added cells did not demonstrate any free-floating spheres but instead developed into large adherent spheres that flattened into monolayers and extended many neurites, whereas cells exposed to excess CSPG remained mostly rounded as free-floating spheres (Fig. 4A and movies 1-3 for mouse, Fig. S5A and movies 4-6 for chick). The highly conserved nature of the hindbrain was further emphasized by the corresponding patterns observed in primary cultures of both chick and mouse embryos. Quantification of neurite length and eccentricity in the cultures illustrated the contrasting effect of modifications in CSPG levels (Fig. 4C,D). Eccentricity can range from 0- 1; 0 represent a full-rounded sphere, while neurite extending from the growing spheres and formation of monolayers consequently increase the eccentricity value. As expected, eccentricity was highest in the ChABC-treated cells and lowest in the excess CSPG group (Fig. 4C,D). For both species, neurite length increased upon treatment with ChABC, simulating the enhanced formation of neurites, hence indicating a more differentiated cell state. Immunostaining the cultures for Tuj1 further highlighted the extensive neurite formation upon CSPG loss, comparing to both control and excess CSPG-treated cells, as the latter did not display nearly any Tuj1^+^ neurites (Fig. 4B for chick, Fig. S5B for mouse). Co-staining the cells for Sox2/Tuj1 further demonstrated this phenotype, as Sox2 levels decreased along with expanded staining of Tuj1 in neurites of the CSPG-depleted cultures. (Fig. 4E). Altogether, the in vitro results recapitulated the in vivo outcomes, collectively suggesting a conserved role of CSPG in promoting a non-differentiated state in cells in the hindbrain.

**Figure 4.**
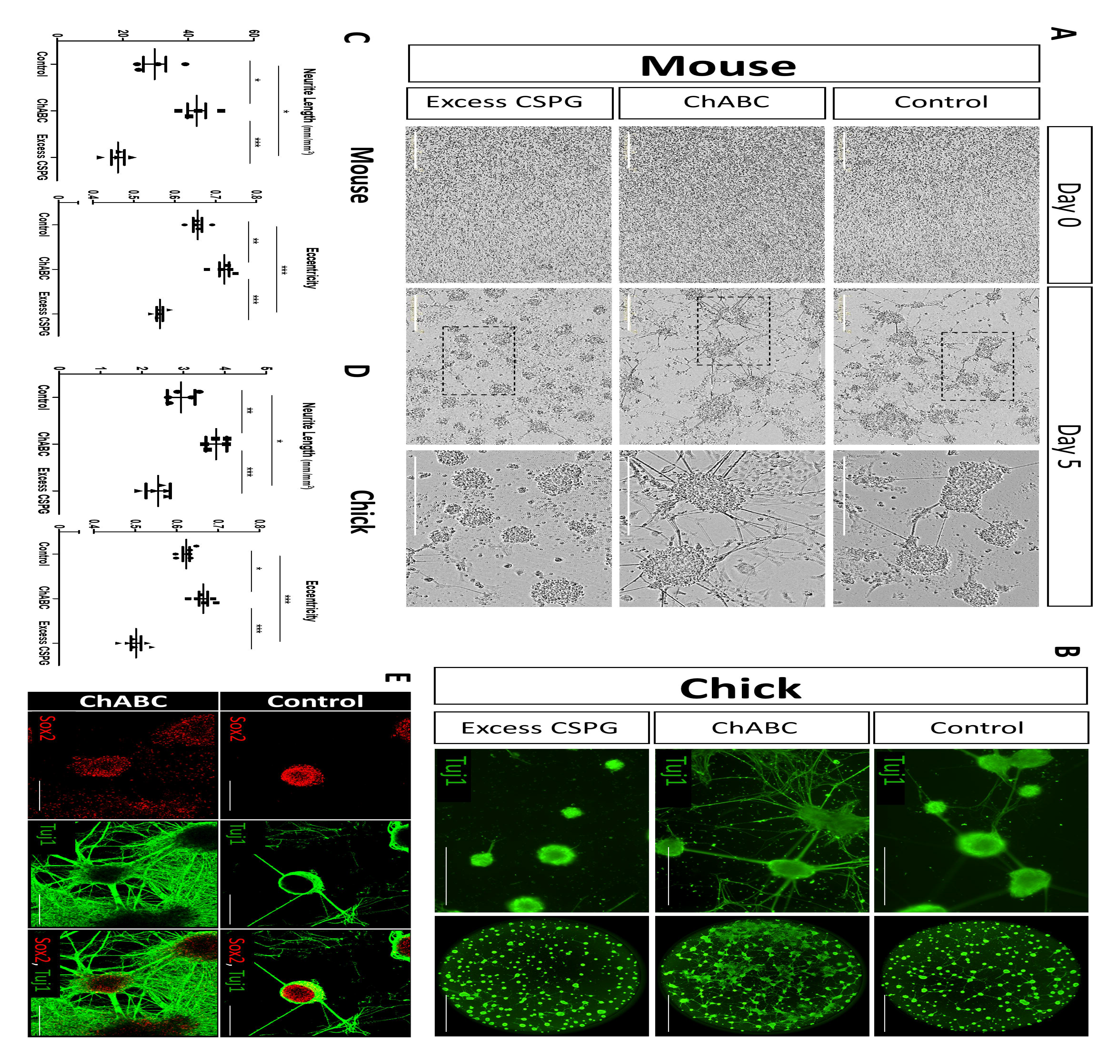
Modifications in CSPG levels in vitro induces shifts in hindbrain cells’ behavior. **(A)**: Representative phase images from time-lapse analysis of primary cell cultures of E10.5 mouse hindbrains on day 0 and 5 of incubation, treated with BSA (control), ChABC or excess CSPG. Higher-magnification of boxed area are presented on the right. **(B)**: Primary cell cultures of st.18 chick hindbrains on day 5 of incubation immunostained for Tuj1 following treatment with ChABC or excess CSPG. Whole-well images of the Tuj1-stained cells presented on the right. (**C,D**): Quantification of eccentricity and neurite-length at day 5 of incubation in mouse and chick cell-cultures. Each dot represents an average of 6 wells from 4 experimental replicates for mouse, or 6-8 wells from 5 experimental replicates for chick. **(E):** Confocal images of primary cultures of st.18 chick hindbrains on day 5 of incubation following treatment with ChABC, immunostained for Tuj1 and Sox2. Data are mean±s.d. (one-way ANOVA with a post-hoc Tukey test).* P< 0.05 ** P< 0.005 *** P<0.0005. Scale bar in A,B= 400 µm, 2000 µm for whole-well images, in E= 100 µm.

### Transcriptomic profiling of CSPG-based separated hindbrain cells

Since CSPG is highly expressed in the HBs, particularly surrounding Sox2+ cells, we employed its membrane-bound expression (Fig. 1, Fig. S1) to isolate the HB^Sox2^ cells, aiming to perform a comparative bulk RNA-seq analysis to fully elucidate the HB’s transcriptome. ∼30 hindbrains of st.18 chick embryos were pooled, dissociated into single cells, immuno-stained for CSPG and underwent FACS (Fig. 5A). This sorting yielded two cell fractions; CSPG-expressing cells (CSPG^+^), which were expected to be enriched in HB cells and comprised 18.4% of all cells, and CSPG-negative cells (CSPG^-^), which made up 53.5% of all cells and consisted mostly of Rh cells, as well as the mantle layer of the HBs (Sox2-). Cells that displayed low levels of CSPG expression were excluded from the analysis (Fig. S6). Notably, DAPI^+^ dead cells were also omitted from the gating, resulting in 95.8% of total hindbrain cells that were used for each FACS series. This procedure was repeated six times. All biological replicates were forwarded for RNA-seq.

**Figure 5.**
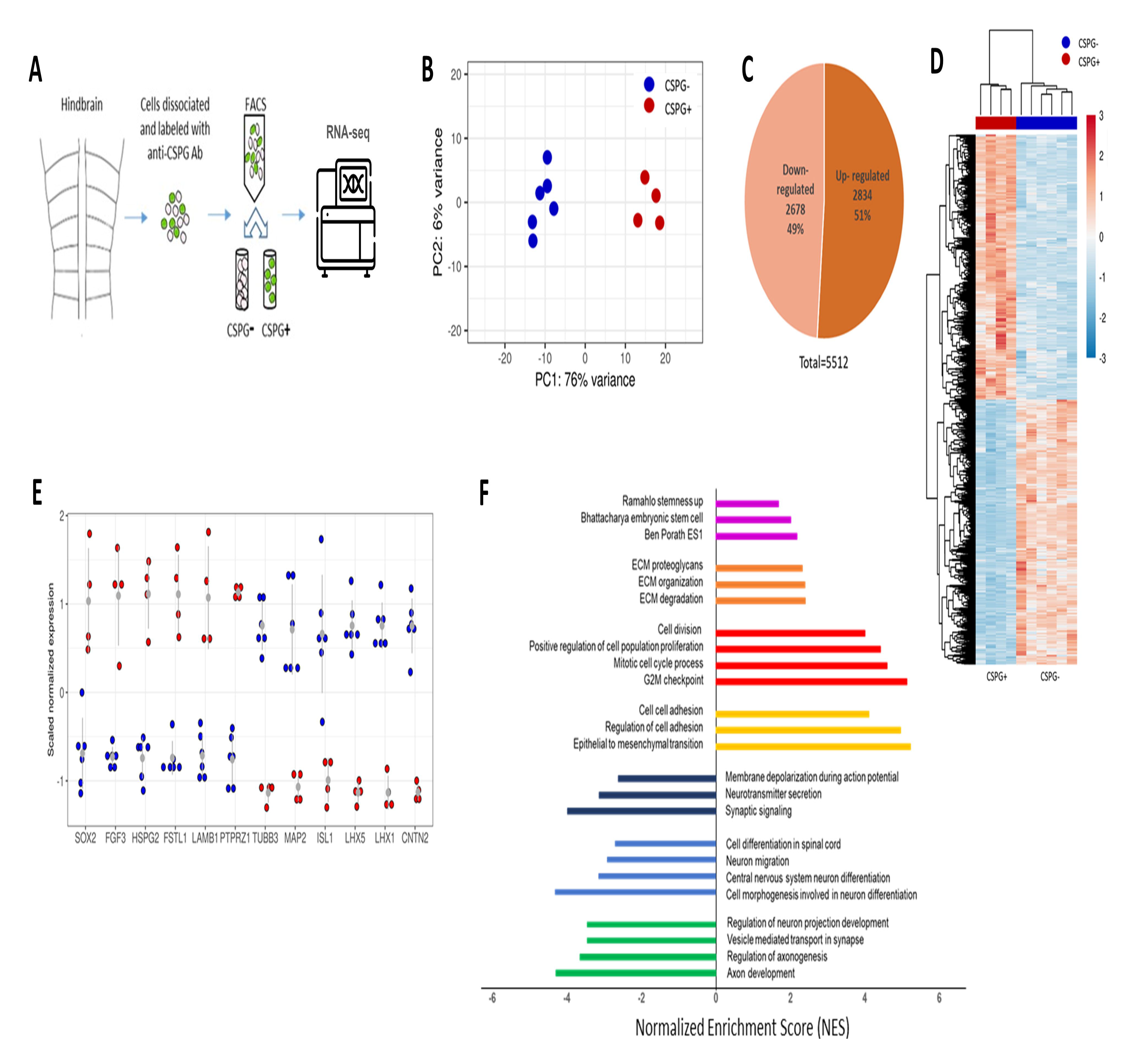
Differential gene expression and genetic pathways distinguish CSPG+ from CSPG-cell groups. **(A):** Illustration of the experimental procedure. **(B):** PCA based on the overall expression pattern of the 6 CSPG- (blue) and 4 CSPG+ (red) samples, showing the first two components and their percentage of the total variance. **(C)**: Distribution of up and down-regulated genes. Genes with a padj<0.05 were included. **(D)**: Heatmap representation of scaled normalized expression signals. Values are colored according to the scale on the right (blue= CSPG-values; red= CSPG+ values). Genes (rows) and samples (columns) were hierarchically clustered as shown by the dendrograms to the left and above the heatmap, respectively. Sample group identity is shown through text below and color annotation above the heatmap. **(E):** Scaled normalized signal of specific genes shown for each sample, colored by group identity (blue= CSPG-; red= CSPG+). Grey dots represents average signal, lines indicating standard deviation. **(F):** Gene-set enrichment analysis showing upregulated (purple, orange, red, yellow) and down-regulated (blue, light blue, green) gene-sets in the CSPG+ group. Change degree is measured by the normalized enrichment score (NES).

Principal component analysis (PCA) of the expression signal showed that the CSPG^+^ and CSPG^-^ replicates were clustered into two highly distinct groups (Fig. 5B). Differential gene expression (DEG) analysis revealed 5512 differentially-expressed genes (adjusted p value < 0.05), out of which 2834 were significantly upregulated and 2678 were significantly downregulated in CSPG^+^ cells compared to the CSPG^-^ group (Fig. 5C), providing a rich data source for further analyses. The expression signal of these differentially expressed genes was reproducible between our biological replicates, as can be seen by the unsupervised hierarchical clustering of samples (Fig. 5D), further indicating the robustness of our experimental design. RNA-seq results for specific genes, selected based on previous knowledge of their expression patterns, are presented in Fig. 5E. For instance, *SOX2*, *FGF3*, *FOLLISTATIN* (*FSTL1*), *LAMB1*, *HSPG* and *PTPRZ1* (a membrane-bound type of CSPG), all previously shown to be enriched at chick HB cells (Heyman et al., 1995; Peretz et al., 2016; Weisinger et al., 2012), were significantly upregulated in the CSPG^+^ group. In contrast, various types of neural differentiation markers, previously reported to be expressed along the mantle layer of the chick hindbrain, such as class III beta-tubulin (*TUBB3*), *MAP2*, *ISL1*, *LHX1*, *LHX5*, and *CNTN2* (also known as *TAG1*) (Kohl et al., 2012; Peretz et al., 2016; Peretz et al., 2018), were significantly upregulated in the CSPG^-^ cell fraction. These results confirm the separation of the hindbrain cells into the expected cell groups.

Gene-set enrichment analysis (GSEA) of the RNA-seq data revealed significant variances in the expression of genes from multiple functional categories between the groups (Fig. 5F). Gene-sets related to embryonic stem cells and cell division were upregulated in the CSPG^+^ cells (Fig. 5F, purple/red bars), along with gene-sets related to cell adhesion and ECM organization (Fig. 5F, yellow/orange bars). Conversely, multiple gene-sets linked to neural differentiation, axonal projection and neuronal activity were upregulated in the CSPG^-^ cell group (Fig. 5F, green/light blue/dark blue bars). Overall, it appears that the CSPG^+^-HB cells display typical characteristics of amplifying NPSCs with a well-defined ECM, whereas the CSPG^-^ cells embody characteristics of differentiating and mature neuronal cells. Moreover, the GSEA results are consistent with our previous findings, where localization of Sox2 and other progenitor markers was confined to the ECM-rich HBs, while neuronal differentiation occurred in the Rhs and/or mantle zone of the HBs (Peretz et al., 2016).

To further investigate some of the most enriched gene-sets of the GSEA analysis (Subramanian et al., 2005), expression of the full sets of genes was presented as hierarchically clustered heatmaps. This showed that a high proportion of the genes in these sets are clustered with a distinct expression pattern of either up or down-regulation. Using the embryonic stem cell pathway 'Ben Porath ES1', multiple genes related to maintenance and self-renewal of stem cells, which were upregulated in the CSPG^+^ group, clustered together in a dense area of the heatmap (Fig. 6A). For instance, the transmembrane glycoprotein *PROM1*, the zinc finger TFs *SALL1*&*4*, the ETs-related TFs *ETV1*&*5* and the cell cycle regulatory genes *CDK1* and *CDC20* (Akagi et al., 2015; Exner et al., 2017; Walker et al., 2013; Wei et al., 2021; Yamano, 2019), all displaying remarkable upregulation. Genes related to cancer stem cells and tumorigenesis, such as *KPNA2*, *ECT2*, *ERCC6L*, *TUBB4B* and the cell-cycle regulators *DLGAP5*, *KIF2C* and *BUB3*, (Christiansen and Dyrskjøt, 2013; Dharmapal et al., 2021; Gong et al., 2020; Pu et al., 2017; Silva and Bousbaa, 2022; Sun et al., 2017; Tsou et al., 2003), were also significantly upregulated in the CSPG^+^ group, and clustered well within the same area. Notably, *SOX2* and *PTPRZ1* were also found in this gene set, indicating its relevance to HBs. Concurrently, even though not contributing to the gene set enrichment score, evaluation of the most notable down-regulated genes in this gene-set further confirmed the NPSC-properties of the CSPG+ cells. Genes involved in neural differentiation, axonogenesis and neuroectoderm induction such as *ROBO1*, *OLFM1*, *CRMP1*, *KIF5C*, *ADD2* and the TFs *ZIC2*&*3* (Aruga, 2004; Guthrie, 2004; Kanai et al., 2000; Matsuoka et al., 1998; Nakaya et al., 2008; Yamashita and Goshima, 2012), were notably downregulated in the CSPG^+^ group (Fig. 6A).

**Figure 6.**
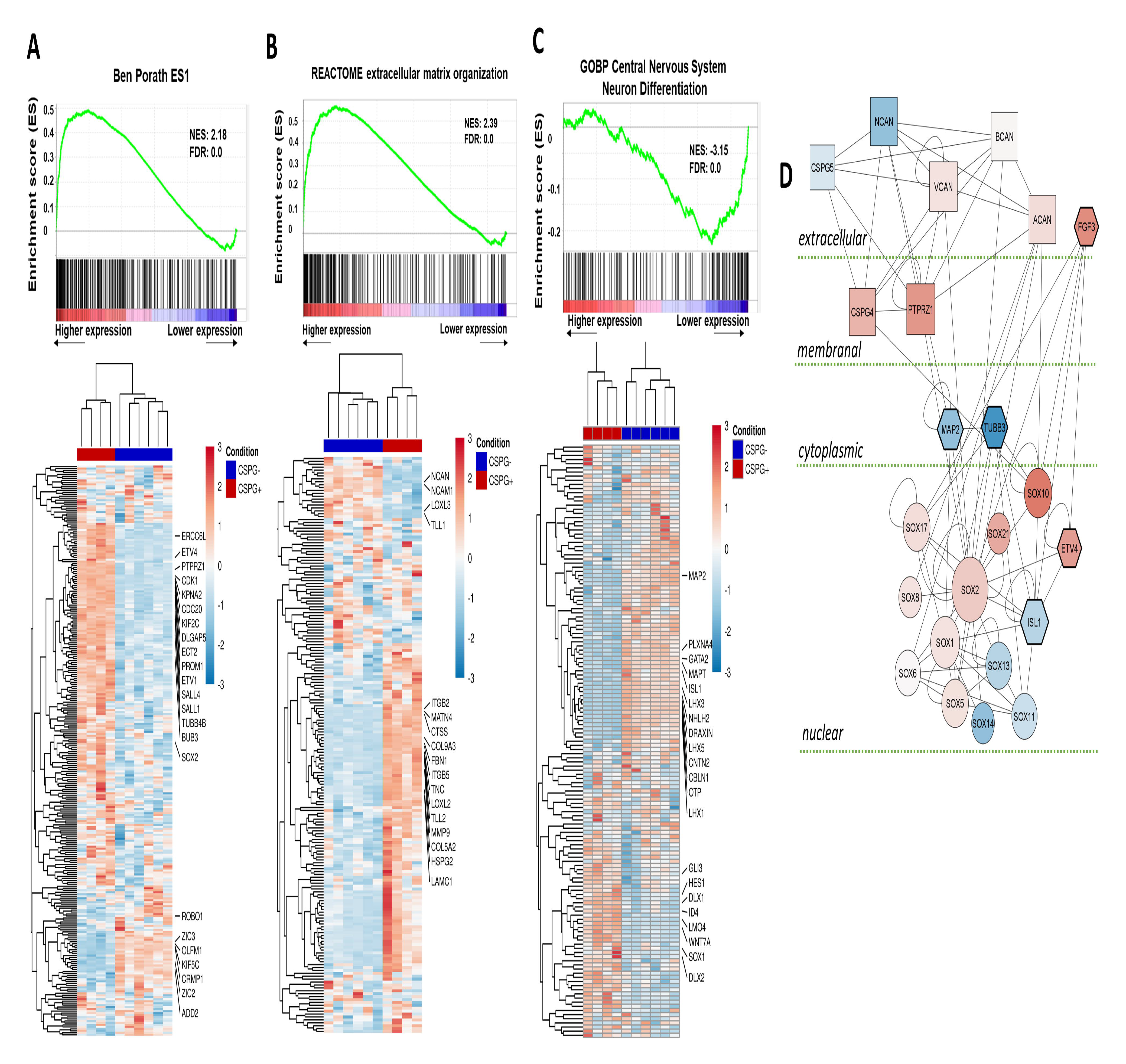
Gene set enrichment analysis of selected pathways. **(A-C):** At the upper part, GSEA-generated graphics showing the distribution of genes from the stated gene-set, represented by vertical lines, at either the left side (A and B, upregulated in CSPG+ groups) or at the right side (C, down-regulated in CSPG+ groups). Heatmap of genes from stated gene-set is presented below. (**D):** A protein-protein network of functionally important genes in CSPG+ vs CSPG-groups. Each shape represents a protein, lines are connecting between interacting proteins (details in Methods). Blue: down-regulated genes. Red: upregulated genes. Round: Sox genes. Square: CSPGs. Hexagon: experimentally validated genes.

Furthermore, analysis of the 'Reactome ECM Organization' gene-set revealed many ECM-related genes that were markedly upregulated in the CSPG^+^ cell fraction (Fig. 6B). These included the cell surface integrin proteins *ITGB2*&*5* (Gardiner, 2011; Ikeshima-Kataoka et al., 2022), various ECM proteins, such as the fibrillar proteins fibrillin (*FBN1*) and collagen sub-types *COL9A3*&*5A2* (Ricard-Blum, 2011), the ECM-adaptor protein matrilin (*MLN4*) (Uckelmann et al., 2016), the tenascin glycoprotein family member *TNC* (Midwood et al., 2016), the sulphated proteoglycan *HSPG2* (Sarrazin et al., 2011), the laminin glycoprotein member *LAMC1* (Aumailley, 2013), and ECM-remodeling proteases, such as *MMP9* (Monsonego-Ornan et al., 2012) and Cathepsin C (*CTCC*) (Tran and Silver, 2021). Conversely, ECM and cell-adhesion proteins that participate in axonal growth and neural differentiation, such as *NCAM* (Paratcha et al., 2003) and the CSPG-soluble protein *NCAN* (Zhou et al., 2001), were markedly downregulated in the CSPG^+^ group, further illuminating the non-differentiated state of this group.

Synchronously, many genes related to neural specification, differentiation, migration and axonogenesis have been significantly upregulated in the CSPG^-^ group, contributing to the enrichment of the 'GOBP Central Nervous System Differentiation' gene-set (Fig. 6C). These included several types of TFs, such as the ventral neuronal markers *GATA2* and *ISL1* (Liang et al., 2011; Zhou et al., 2000), the dorsal interneuron markers *LHX1/3/5* (Hirsch et al., 2021; Pillai et al., 2007), the dopaminergic neuronal marker *OTP* (Ryu et al., 2007) and the regulator of precerebellar nuclei migration *NHLH2* (Schmid et al., 2007). Moreover, various axonal-growth cues and receptors, like the semaphorin receptor *PLXNA4* (Suto et al., 2005), the repulsive signal *DRAXIN* (Ahmed et al., 2011), the chemoattractant signal *CBLN1* (Han et al., 2022), the microtubule-associated proteins *MAP2* and *MAPT* (Riederer and Matus, 1985) and the cell adhesion molecule *CNTN*, which promotes axon guidance and fasciculation (Stoeckli and Landmesser, 1995), were all significantly upregulated in the CSPG^-^ cell group (Fig. 6C). Accordingly, factors that promote cell proliferation and prevent neural differentiation in NPSCs/neuroepithelial cells were clearly downregulated in the CSPG^-^ group. These included the TFs *HES1* (Kageyama et al., 2008), *DLX1*&*2* (Carrillo-García et al., 2010), *ID4* (Bedford et al., 2005), *LMO4* (Kashani et al., 2006), *SOX1* (Venere et al., 2012), and the cell-cycle-regulated protein *HURP* (Hepatoma Up-Regulated Protein) (Tsou et al., 2003).

Finally, protein-protein interaction network analysis revealed the putative cross-talks between different CSPG subtypes, FGF signaling components, neural differentiation markers and various Sox genes that are up or down-regulated in the CSPG^+^ cell group, in accordance with their extra or sub-cellular localization (Fig. 6D). This analysis further demonstrates the direct interaction of Sox2 with the membranal CSPG subtype PTPRZ1 and the soluble signal molecule FGF3, in agreement with their spatial expression patterns (Figs 1,2) (Weisinger et al., 2012). Collectively, the divergence of the whole-transcriptomic data together with the enrichment of the gene-set categories and networks, substantiate that HB cells are a sub-population of NPSCs which aggregate in particular niches at the HBs, differing from the adjacent more differentiated CSPG-/Rhs cells.

### CSPG+ HB cells reveal typical NPSC-behavior in vivo

As the transcriptome of HB^Sox2^ cells appeared to resemble that of other types of NPSCs, we next examined whether these molecular properties are coupled with typical NPSCs’ behavior. To separate the HB cells from the rest of hindbrain cell population, we once again relied on the membrane-bound CSPG expression in HB^Sox2^ cells (Fig. S1). Hindbrains of chick embryos were dissociated and immuno-labeled for CSPG, then passed through a magnetic-based immuno-column, ultimately providing two viable CSPG+/CSPG-cell fractions (Miltenyi et al., 1990) (Fig. 7C). This sorting method was selected this time, as the cells demonstrated a higher viability during prolonged incubation, in comparison to FACS sorted cells. Flow-cytometry analysis confirmed an efficient separation, which demonstrated a 4-fold increase in cells expressing CSPG in the CSPG^+^ fraction compared to the CSPG^-^ fraction (Fig. S7). The CSPG^+^ and CSPG^-^ cell groups were seeded and live-imaged for 5 days, followed by immunostaining for Tuj1 at the end of the incubation. Noticeably, the two cell fractions displayed distinctly dissimilar characteristics; CSPG^+^ cells mostly gathered into rounded spheres with very few cells that adhered and extended neurites, whereas the CSPG^-^ cells largely adhered and showed high extension of neurites (Fig. 7A; Movie 7 for CSPG^+^; Movie 8 for CSPG^-^). This extensive neurite formation was further demonstrated by a whole-well view of Tuj1-immunostained cells, as the CSPG^-^ cells revealed a substantial formation of entwined network of Tuj1-expressing neurites, extending in and between the adhered spheres, while the CSPG^+^ cells were found in rounded spheres, almost completely devoid of Tuj1+ neurites (Fig. 7B). As monolayer structure is typical for differentiating neurons, formation of monolayer was quantified in the two cell fractions and was found to be higher in the CSPG^-^ group (Fig. 7D). Neurite length and eccentricity were also significantly higher in those cells, further revealing the more differentiated nature of the CSPG^-^ cells compared to the CSPG^+^ population (Fig. 7E,F). Together, the noticeably distinguished behavior of the two cell populations further verifies that the CSPG^+^ fraction is enriched with cells displaying typical NPSC-phenotypes, whereas the CSPG^-^ fraction is comprised of more differentiated hindbrain cells.

**Figure 7.**
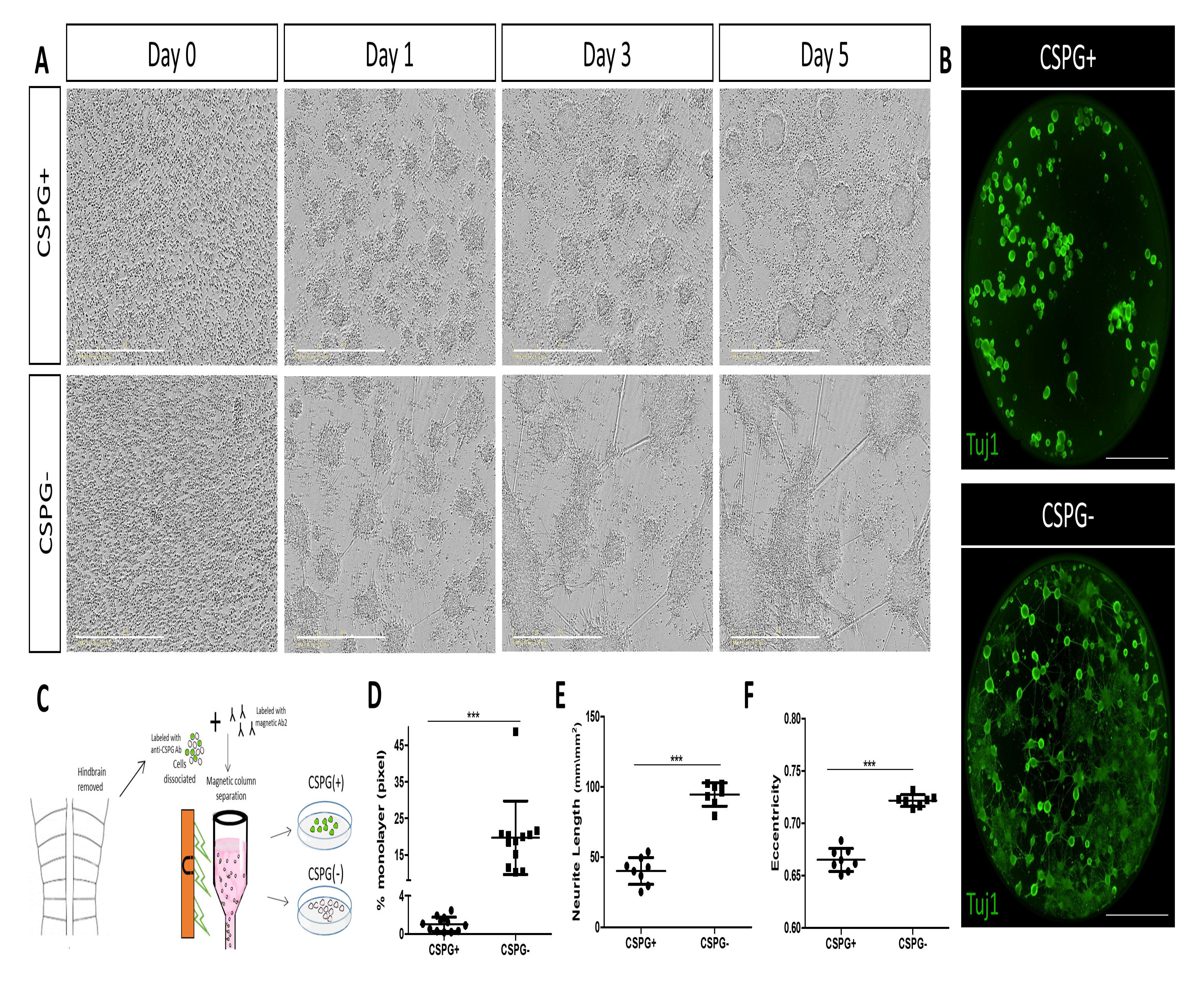
CSPG-based separated cells show distinct cell characteristics in vitro. **(A):** Representative phase images from time-lapse analysis of st.18 chick hindbrains cell cultures separated for CSPG+ and CSPG-groups. **(B):** Whole-well imaging of Tuj1-stained CSPG+ and CSPG-cells. **(C):** Illustration of the magnetic-based immuno-column separation of hindbrain cells. (**D-F):** Quantification of monolayer, neurite length and eccentricity in CSPG+ vs. CSPG-cells. Each dot represents an average calculated in one well at day 5 of incubation. n=8 wells for each group from 3 experimental replicates. Data are mean±s.d. (one-way ANOVA with a post-hoc Tukey test). *** P<0.0005. Scale bar in A= 400 µm, in B= 2000 µm.

### CSPG is required to maintain HB^Sox2^ cells in their NPSC state

Finally, to fully uncover the effect of CSPG on HB^Sox2^ cells, we sought to directly examine the behavior of the HB/CSPG^+^ cells following CSPG removal. Live-imaging analysis showed the untreated CSPG^+^ cells typically aggregating to form rounded spheres (Fig. 8A, Movie 9). However, addition of ChABC to the CSPG^+^ cells caused them to rapidly adhere and extend many neurites (Fig. 8A, Movie 10), thus exhibiting a remarkable resemblance to the CSPG^-^ cell fraction (Fig. 7A). To quantify the shift in the cells’ state, formation of monolayers and development to adhered/floating type of spheres were measured in the control and ChABC-treated CSPG^+^ cells (Fig. 8D-F). Appropriately, control cells formed almost exclusively floating spheres, while formation of monolayers was highest in the ChABC-treated group. This behavior was comparable to that detected in the CSPG^-^ cell group, which also generated mostly adherent spheres, indicating that loss of CSPG in HB^Sox2^ cells is sufficient to shift them into a more differentiated state, same as evident in CSPG^-^ cells. Immunostaining with Sox2 and the neural differentiation marker Map2 further elucidated the change in the cell differentiation state, as Sox2 expression was found to be more profound in control cells compared to a weaker expression in the ChABC-treated cells (Fig. 8B). Moreover, Map2, which was expressed in the outer layer of the spheres in the control and ChABC-treated cells, was also found in the extending neurites, formed exclusively in the latter group (Fig. 8B). Real-time RT PCR analysis of relative gene expression in the two cell groups showed a decrease in *Sox2* and an increase in *Map2* levels in CSPG^+^ cells upon treatment with ChABC, further confirming the shift in the differentiated state upon loss of CSPG (Fig. 8C).

**Figure 8.**
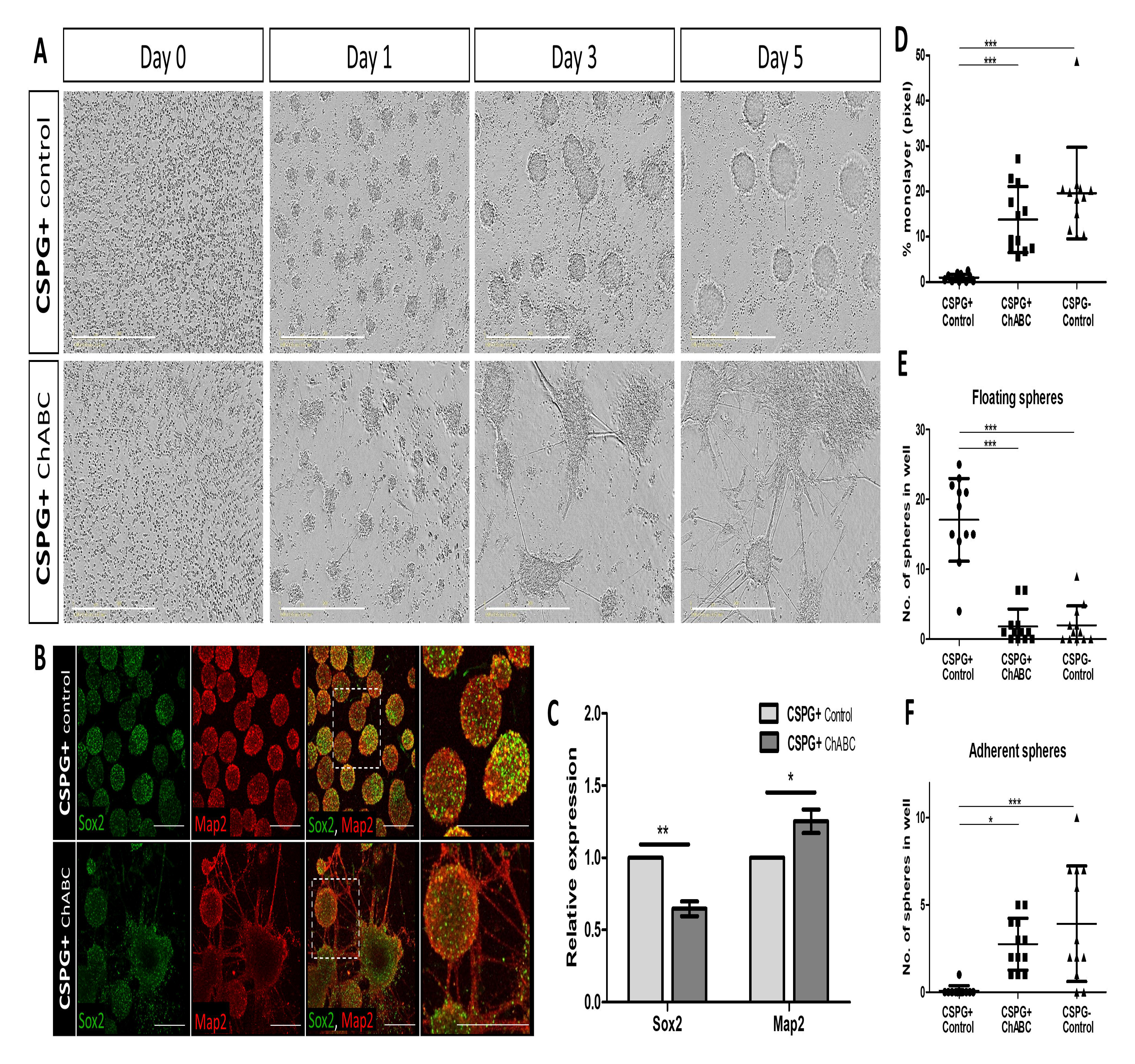
CSPG loss promotes HB’s cell differentiation. **(A):** Representative phase images from time-lapse analysis of CSPG+ control or ChABC-treated cells. **(B):** Confocal images of control and ChABC-treated CSPG+ cells on day 5 of incubation immunostained for Sox2 and Map2. Higher-magnification of boxed are presented to their right. **(C):** Real-time RT PCR analysis of *Sox2* and *Map2* expression in control and ChABC-treated CSPG+ cells. Data are mean±s.d. from 3 experimental replicates; n=4 wells for each group (two-tailed unpaired t-test). **(D-F):** Quantification of monolayers and floating/adherent spheres in control CSPG+ and CSPG-cells and in ChABC-treated CSPG+ cells. Each dot represents an average calculated in one well at day 5 of incubation. n=12 wells for each group from 3 experimental replicates. Data are mean±s.d. (one-way ANOVA with a post-hoc Tukey test). * P< 0.05 ** P< 0.005 *** P<0.0005. Scale bar in A= 400 µm, in B= 100 µm.

## DISCUSSION

This study reveals that HBs are enriched with cells expressing Sox2 and CSPG in mouse and chick embryos and unveiled the function of CSPG in maintaining these cells as NPSCs, as they reside in designated niches in-between the differentiating Rhs. The transcriptomic profile of HB cells uncovered a substantial number of DEGs that participate in multiple types of embryonic/neural/cancer stem cell-related pathways, and in ECM-enriched pathways, which were not identified in HB cells before. Correspondingly, neural differentiation-related genes and pathways were found to be upregulated in the non-HB’s cells. Ultimately, this study highlights the unique position of HBs in amniotes as CSPG-enriched niches of NPSCs in-between the differentiating Rhs, as well as contributes new data regarding genes and pathways active in these cells, which emphasizes the significance of compartment boundaries during development.

### HB cells are NPSCs

Highlighting the uniqueness of HBs as repetitive CSPG-rich niches of NPSCs in the hindbrain, raises the question of how similar are they to other NPSC’s types. Comparing our transcriptome data with knowledge on NPSCs from other domains reveals that in addition to shared expression of Sox2 among many types of NPSCs, other NPSC landmark genes are also expressed in the HBs. This includes, for example, the membranal protein prominin-1 (*PROM1* or *CD133*) and the glutamate transporter family member *GLAST1* (*SLC1A3*). These markers are paramount for NSCs from the SVZ and DG to self-renew and to acquire multi-lineage differentiation capacities (Coskun et al., 2008; Lee et al., 2005; Uchida et al., 2000). Interestingly, PROM1 and GLAST1 are also upregulated in brain cancer cells and correlate with poor prognosis, demonstrating the intricate relationship between NSCs and brain tumors (Reya et al., 2001; Singh et al., 2004). These findings raise the question as to whether abnormal development/maintenance of HB cells may participate in brainstem tumors.

The milieu surrounding NPSCs in various CNS domains is enriched with different ECM proteins that dictate their proliferation, differentiation and fate (Kazanis and ffrench-Constant, 2011; Preston and Sherman, 2011; Walma and Yamada, 2020). For instance, the secreted glycoprotein Tenascin-C (TNC) and the ECM receptor integrin b (ITGB) modulate such processes in telencephalic or spinal-cord NPSCs, as their loss-of-function decreased the number of NPSCs (Faissner et al., 2017; Garcion et al., 2001; Karus et al., 2011; Temple, 2001; Theocharidis et al., 2021). Our transcriptomic analysis found that *TNC* and *ITGB2/ITGB5*, were upregulated in HB cells, suggesting similar role in promoting the development of the HB-NPSCs. Notably, in chick mesencephalic neural progenitors, integrin was found to activate Wnt7A signaling, which in-turn induces the expression of the ECM molecule Decorin (DCN), that promotes neurogenesis in neighboring cells (Long et al., 2016). Intriguingly, *WNT7A* and *DCN* were also upregulated in the HB/CSPG^+^ cell group, implying a conserved function for integrins in midbrain and hindbrain progenitors.

Several cell-cycle regulatory genes have been identified in various types of NPSCs and cancer-stem cells. For example, the kinesin molecule Kif2C and the anaphase-promoting factor CDC20 regulate proliferation of NSCs or tumorigenic brain cells (Li et al., 2022; Qi et al., 2020; Sun et al., 2017; Tu et al., 2022; Xie et al., 2015; Yang et al., 2019). These genes were upregulated in HB cells, along with the DNA excision repair gene *ERCC6L*, a pan-cancer marker that promotes growth and invasion in various cancers including glioma (Chen et al., 2022; Xie et al., 2019). Mutations in this gene are linked to Cockayne syndrome, a rare inherited disorder characterized by neural abnormalities (S. Wang et al., 2020). This gene, which may be involved in regulating the proliferation of HB cells, has not yet been reported in other types NSCs.

Altogether, these examples reveal several emerging genetic similarities between HBs’ NPSCs and other NPSCs. The outcomes of their mis-regulation in other contexts could possibly unravel their role in regulation of the HB cells as NPSCs.

### The role of CSPS in the HBs

CSPG has been previously found to regulate NPSC’s state in the brain. Yet, its removal from NPSCs using ChABC have reported different outcomes, with some reporting reduced neurogenesis and proliferation and others reporting increased proliferation, differentiation and migration (Gu et al., 2009; Sirko et al., 2007; Yamada et al., 2018). Our transcriptomic profiling reveled that *PTPRZ,* a transmembrane protein of the CSPG family, was significantly elevated in HB cells. This correlates with the membranal-bound pattern of CSPG observed in our immunostaining, altogether suggesting that the HBs are enriched with PTPRZ. Previous studies using cortical and telencephalic NSCs have reported PTPRZ expression and found that its degradation using ChABC decreased their self-renewal and neurosphere formation and increased their differentiation in vitro (Sirko et al., 2007; Sirko et al., 2010b; von Holst et al., 2006). Our data further emphasizes the significance of this CSPG in hindbrain NPSCs, as treatment with ChABC decreased Sox2 expression and promoted cell differentiation in vivo and in vitro. Notably, the effect of CSPG modifications on NSC’s fate differs in different CNS domains; interference with the sulfated state of the glycosaminoglycan (GAG) chains caused cortical-derived NSCs to differentiate towards the astrocytic lineage, while spinal cord NPSCs generated more immature neurons in vitro (Karus et al., 2012; Sirko et al., 2007). Further investigation is required to determine the fate of NPSCs in the HBs, specifically when it is either enriched or depleted of CSPG.

How can CSPG maintain the HB cells undifferentiated? As a highly abundant ECM factor in the CNS, CSPG was found to interact with multiple signaling molecules, neurotrophic factors, ECM components and cell-adhesion proteins (Djerbal et al., 2017). Yet, most of these interactions occur in processes that follow NPSC’s stages, including neuronal migration, axonal pathfinding and synaptogenesis (Dyck and Karimi-Abdolrezaee, 2015; Mencio et al., 2021; Miller and Hsieh-Wilson, 2015; Rogers et al., 2011). Interestingly, CSPG is a key inhibitor of axonal growth upon CNS injury, hence, its elimination is essential for regeneration (Brown et al., 2012; Griffith et al., 2017). Modified CSPG levels are also related to mental and neurodegenerative disorders (Jang et al., 2020; Schultz et al., 2014), signifying the multi-faceted roles of CSPG in the CNS (Zhang and Chi, 2021). Contrary to its function in these processes, its role in NPSCs is less known. Several studies have reported that CSPG is required for maintaining a stemness state in NPSCs in the SVZ, and have suggested that this activity is mediated by interaction with FGF-2 to promote cell proliferation (Bian et al., 2011; Roll et al., 2022; Sirko et al., 2010a). In spinal cord NSPCs, such FGF-2 activity was recently found to depend on the sulfation patterns of the CSPG-GAG chains that operate as docking sites for specific proteins (Schaberg et al., 2021). Interestingly, our RNA-seq data did not find a differential expression of *FGF-2* in the two CSPG-separated cell groups. However, other FGFs, including *FGF-3,-8,-10,-18,-22*, were upregulated in the HB-cell fraction, suggesting that local CSPG may prevent cells from undergoing differentiation by acting as a co-receptor for these FGFs. This possibility is supported by the upregulation of several FGF-downstream target genes in the HB cells, (such as *ETV-4,-5*), as well as by our previous findings on the presence of di-phosphorylated ERK (dpERK) in these domains (Weisinger et al., 2012). Yet, it is possible that other signals, receptors or ECM factors interact with CSPG at the HBs (Tham et al., 2010). For instance, transcripts of various ECM-related factors, (collagens, integrins, laminin, fibronectin), which have been reported to interact with CSPG in other processes, such as metastasis, CNS injury, and axonogenesis (Avram et al., 2014; Knutson et al., 1996; Ohtake and Li, 2015), were upregulated in the HB’s RNA-seq. Upon illuminating the significance of CSPG in regulating the state of HB cells, deciphering the pathways and the participating factors should be the next step.

### HBs in different model systems

In contrast to the limited data on HBs in amniotes, extensive research has been performed in zebrafish. Zebrafish HB cells were found to act as organizing centers to induce neurogenesis in rhombomere-flanking zones and to repel axons and drive their accumulation in the Rhs (Gonzalez-Quevedo et al., 2010; Terriente et al., 2012). In agreement with our previous finding on chick HBs as reservoirs of NPSCs (Peretz et al., 2016), subsequent works confirmed that in zebrafish HBs similarly serve as pools of progenitors in an active proliferative state, regulated by Yap/Taz-TEAD activity (Voltes et al., 2019). Recent monitoring of the spatiotemporal dynamics of HB cells in zebrafish have revealed that they initiate as neuroepithelial-stem cells that divide symmetrically, while they later shift to become radial-glia progenitors undergoing asymmetrical division to contribute neurons to the hindbrain (Hevia et al., 2022). This transition was found to be triggered by Notch3 signaling. Intriguingly, while *NOTCH3* was not detected in our transcriptomic analysis, *NOTCH2* is upregulated in the chick HB cells, whereas two delta ligands (*DLL1,4*) were upregulated in the non-HB fraction. These findings may indicate a conserved lateral-inhibition manner of action of HB cells in avian and teleost to preserve HB cells undifferentiated. This is further supported by the upregulation of the Notch downstream target *Hes1* in the HB’s transcriptome, as also previously found in mice HBs (Baek et al., 2006). Yet, the Notch-signaling supporting factors, radical fringe (*rfng*) and lunatic fringe (*lfng*), which are expressed in zebrafish HB cells and prevent them from undergoing differentiation (Nikolaou et al., 2009; Voltes et al., 2019), were not detected in our RNA-seq, nor in previous studies in mice (Moran et al., 2009). These differences may indicate for inter-species variations in the of HB’s gene profile, consistent with other factors that are expressed in HBs of avian and mammalian but not in fish, such as FGF3 and Sox2. These species-specific properties are also emphasized by the fact that in zebrafish Rh centers are another non-neurogenic zone that also regulate neural differentiation in their neighboring domains(Cheng et al., 2004; Gonzalez-Quevedo et al., 2010). This leads to the formation of repetitive neurogenic stripes in the Rhs, found in-between the HBs and Rh centers, which are not evident in amniotes.

Further comparison between HBs of zebrafish and chick can be drawn from a single-cell RNA-seq done in zebrafish hindbrain (Tambalo et al., 2020). In this analysis, three main cell clusters were detected: HB cells, Rh center cells and neurogenic cells. The HB’s cluster expressed genes such as *rasgef1b*, *rac3b*, *prdm8*, *follistatin* and *gsk1*. Looking into our transcriptomic data, those genes were either found to not differ between the CSPG^+^/CSPG^-^ groups, or to be upregulated in the non-HB fraction. However, some of their homologues, such as *RASGEF1A*, *RAC1* and *FSTL1*, were upregulated in the chick HB cells. Likewise, *fgf20* and *etv5* transcripts, which have been identified in the Rh centers of zebrafish, were either not detected in the hindbrain *(FGF-20*) or actually upregulated in the HBs (*ETV5*). Nevertheless, the presence of several other FGFs in chick HB cells, along with the upregulation of several FGF downstream targets (*ETV1-5,-4*, *DUSP6*, *SPRED1*), raises the intriguing possibility that the non-neurogenic function of Rh centers in teleost may have been lost in amniotes, while the HB zones have retained a similar role throughout evolution.

## MATERIALS AND METHODS

### 1. Embryos

#### Chick

Fertile Loman chicken eggs (Gil-Guy Farm, Moshav Orot, Israel) were incubated at 37°C for 72-84 hours until reaching the desired Hamburger Hamilton (HH) developmental stage of st.14 or st.18, as specified. A small hole was made in the shell through which 5 ml of albumin were removed using a syringe. Next, a small window was made in the shell to expose or harvest the embryo for further procedures (Kayam et al., 2013).

#### Mice

Wild type (WT) mice C57BL/6 were purchased from Harlan Laboratories (Rehovot, Israel). Mice were mated and females were examined for a vaginal plug (VP) the following morning, this was considered as embryonic day (E) 0.5. After 10.5 days, females were sacrificed and embryos were taken for further procedures (Kalev-Altman et al., 2020). Mice were kept in the Hebrew University Specific Pathogen Free animal facility according to animal care regulations. All procedures were approved by the Hebrew University Animal Care Committee (license number 18-15452-1).

### 2. Hindbrain cell culture experiments

#### 2.1 Primary cultures of chick hindbrain cells

Hindbrain regions of st.18 HH embryos were dissected in sterile PBS with Penicillin-Streptomycin (Pen-Strep, 1:100; Gibco, USA), then placed in a tube containing human embryonic stem cell medium (hESC; DMEM/F-12 1:1 with 20% KnockOut serum replacement, GlutaMax L-alanyl-L-glutamine (2 mM), non-essential amino acids (0.1 mM; all from Gibco, USA), β-mercaptoethanol (0.1 mM; Sigma-Aldrich, USA), Pen-Strep (1:100) and Fungizone (1:500)). Media was next replaced with 1 ml of TrypLE Express (Gibco, USA) to dissociate the tissue into single cells. Following a manual disassociation by pipetting up and down, TrypLE was neutralized with a 10:1 hESC medium and cells were passed through a 100μm mesh strainer to detach adherent cells. Cells were cultured in hESC media, at density of 1×10^5-6^ cells\ml, seeded in a 48\96 well Nunclon Delta Surface culture plate (Thermo Fisher Scientific, USA) and incubated at 37°C in 5% CO_2_ (Peretz et al., 2016; Peretz et al., 2018). For live imaging, cell plates were imaged every 3-6h in IncuCyte S3 Zoom HD/2CLR time-lapse microscopy system, equipped with x20 Plan Fluorobjective (Sartorius, Germany). Time-lapse movies were generated by capturing phase images for up to 5 days of incubation (X. Wang et al., 2020).

#### 2.2 Primary cultures of mouse hindbrain cells

Cell cultures from E10.5 hindbrains were prepared and imaged as described above, with the following modifications; hindbrains were collected into PBS containing calcium and magnesium (Biological Industries, Israel), and cells were grown in hESC media combined with NeuroCult Proliferation Supplement, added with 20 μg of human recombinant EGF, 10 μg human recombinant bFGF and 10 μg 0.2% heparin solution (all from STEMCELL Technologies, Canada).

#### 2.3 Magnetic bead cell sorting of hindbrain cells

Cell separation was done using MACS^®^ MicroBeads cell separation system (Miltenyi Biotec, Germany) according to the manufacturer’s protocol, with slight adjustments. Briefly, 60 hindbrains of st.18 chick embryos were harvested and disassociated into single cells using collagenase type 4 (200 Units\ml, Worthington 47B9407, USA). Cells were then centrifuged at 600 *g* for 10 min, washed in PBS, and re-centrifuged. Next, Cells were incubated with mouse anti-CSPG antibody (#c8053; Sigma-Aldrich, USA) diluted 1:50 in MACS^®^ BSA Stock Solution and autoMACS^®^ Rinsing Solution (1:20, Miltenyi Biotec, Germany) for 1-2h at RT. Next, cells were centrifuged and washed in PBS twice, then incubated with anti-Mouse IgG micro-bead (1:10 in autoMACS Running Buffer, Miltenyi Biotec, Germany) for 30 min at 4°C. Cells were then washed and moved into MACS^®^ cell separation magnetic columns placed on MACS^®^ iMAG separator, allowing the CSPG+ cells to attach to the column, while the CSPG-fraction passed through and collected. The CSPG^+^ cells were finally eluted from the column by removal of the magnetic field, and collected separately. The separated CSPG+ and CSPG-cells fractions were centrifuged, suspended in hESC medium, plated to generate a culture, and grown and imaged as described above. Validation of the proper separation into CSPG+ and CSPG-cell fractions was done using flow cytometry analysis, as described below.

#### 2.4 Treatments

Inhibition of CSPG through digestion of its CS chains was done using addition of 50 mU/ml Chondroitinase ABC (ChABC; Sigma-Aldrich, USA) diluted in 0.01% BSA (Sigma-Aldrich, USA). Addition of external CSPG was achieved using 50 mg/ml proteoglycan from bovine nasal septum (Sigma-Aldrich, USA), diluted in molecular grade water. Both treatments were added to the culture media every 48h. As controls, cells were treated similarly with 0.01% BSA or water.

### 3. In vivo experiments

#### 3.1 Plasmid electroporation

pcDNA3.1-chABC and pcDNA3.1-GFP plasmids (Muir et al., 2010) were mixed 2:1. For control, pcDNA3.1-GFP plasmid was mixed with molecular grade water in a 2:1 ratio. Plasmids were injected into the hindbrain lumen of st.14 embryos using a pulled glass capillary, as previously described (Kohl et al., 2013). L-bent gold electrodes (1 mm diameter) were placed flanking the hindbrain and an electrical current of 25 V was applied in 5 pulses of 45ms with a pulse interval of 300 ms using ECM 830 electroporator (BTX, USA). Following electroporation, PBS was applied over the embryos, eggs were sealed with parafilm and re-incubated at 37°C for additional 24 h before harvesting.

#### 3.2 ChABC injection

50 mU/ml of ChABC, diluted in 15% Pluronic® F127 thermosensitive hydrogel (Sigma-Aldrich, USA), were injected locally into the hindbrain lumen of st.14 embryos in ovo, using a pulled glass capillary. As control, embryos were treated with 0.01% BSA. Following injection, embryos were re-incubated ON in 37°C, allowing the hydrogel to solidify, hence assuring a prolonged exposure of the area to the substance (Pokhrel et al., 2022; Shriky et al., 2020). Post incubation, embryos were harvested and hindbrains were removed.

### 4. Flow-cytometry

Whole hindbrains dissected from st.18 chick embryos, or 5-days old primary cultures, treated as mentioned above, were incubated in Express TrypLE for 10 min at 37°C, then dissociated manually and neutralized with 1:10 hESC medium. Cells were then fixated in 4% paraformaldehyde solution (PFA; Sigma-Aldrich, USA) for 10 min at RT, centrifuged at 600 *g* for 10 min, washed in PBS for 5min, and centrifuged again. Cells were next incubated in blocking solution (0.2% Triton in PBS and 2% goat serum) for 1 h at RT, followed by incubation of 2 h at RT or overnight at 4°C in 1% BSA with primary antibodies (1:300). Following washes and centrifugation, cells were incubated for 2h at RT in 1% BSA with the appropriate Alexa-Fluor secondary antibody (1:300; Life Technologies, USA) dissolved in 0.5% BSA. Next, cells were centrifuged, washed, and centrifuged again as described above. Finally, cells were suspended in clean PBS and passed through an Accuri C6 Flow Cytometer (BD Biosciences, USA). Flow cytometry analysis was performed using BD Accuri C6 software. For validation of the immuno-magnetic separation according to CSPG-expression levels, CSPG+/CSPG-cells were collected and centrifuged for 3 min at 14,000 g, then subjected to the same procedure. Detection of cell death due to exposure to ChABC was achieved using Annexin V-FITC Early Apoptosis Detection Kit (Cell Signaling, USA), done according to the kit user guide with minor modifications. Briefly, treated and control hindbrains were dissociated and prepared as described above, then resuspended in 300 µl 1x Annexin V Binding Buffer. Next, cells were added with Annexin V-FITC conjugate (1:100) and Propidium Iodide (1:30) and incubated on ice for 10 min. Finally, 100 µl of 1x Annexin V Binding Buffer was added to the cells to terminate the reaction, and cells were taken for FACS analysis.

### 5. Flow-cytometry cell sorting

30 hindbrains of st.18 chick embryos were harvested and disassociated into single cells using collagenase type 4 (200 Units\ml, Worthington 47B9407, USA). Cells were then centrifuged at 600 *g* for 10 min, washed in PBS, then centrifuged again. Next, Cells were incubated with mouse anti-CSPG antibody (#c8053; Sigma-Aldrich, USA) diluted 1:50 in MACS^®^ BSA Stock Solution and autoMACS^®^ Rinsing Solution (1:20, Miltenyi Biotec, Germany) for 75 min at RT. Next, cells were centrifuged, and washed in PBS twice, then incubated with anti-Mouse Alexa-Fluor 488 antibody (1:200, Thermo Fisher Scientific, USA) in autoMACS^®^ Running Buffer (Miltenyi Biotec, Germany) for 30 min at RT. Cells were then washed, centrifuged and resuspended and kept in hESC media ON at 4°C. Next, cells were washed with autoMACS^®^ Running Buffer and stained with DAPI (1:200 in autoMACS^®^ Running Buffer), for 5 min at RT, then washed again. 1×10^7^ cells\ml were passed to FACS tubes and sorted using ARIA III FACS (BD Biosciences, USA) into 1 ml autoMACS^®^ Running Buffer. The gating was set according to size and granularity using FSC and SSC to capture singlets and remove debris. The cut-off for sorting the positive (CSPG+) and negative (CSPG-) cells was based on Alexa-Fluor 488 stained\unstained cells, appropriately, with the exclusion of dead/damaged DAPI+ cells, chosen by manual gating.

### 6. Immunofluorescence

#### 6.1 Embryos

Chick or mouse embryos were harvested at st.18 or E10.5, respectively, cleaned from surrounding membranes and fixed in 4% PFA (Sigma-Aldrich, USA) ON at 4°C. Embryos were next washed with PBS, and incubated in blocking buffer (0.1% Tween20 in PBS (PBT) with 5% goat serum) (Biological Industries, Israel) for 2 h. Next, embryos were incubated ON at 4°C in blocking solution with the following primary antibodies; Rabbit-anti Sox2 (1:400; #ab5603; Millipore, USA), Mouse-anti CSPG (1:80; #c8053; Sigma-Aldrich, USA), Mouse-anti Tuj1 (1:400; #ab14545; Abcam, UK); Mouse-anti Map2 (1:200; #ab15452; Sigma-Aldrich, USA). Following washes, embryos were incubated for 2 h at RT with the following secondary antibodies; Goat anti-mouse Alexa488, Goat anti-mouse Alexa594, Goat anti-rabbit Alexa488, Goat anti-rabbit Alexa594 (1:300; diluted in blocking solution; Life Technologies, USA). Next, Embryos were washed with PBS and incubated for 15 min at RT in PBS with DAPI (1:400; Sigma-Aldrich, USA). Following washes with PBS, hindbrains were finally dissected and flat-mounted on slides with FluoroGel with Tris buffer mounting medium (Electron Microscopy Science, USA) (Kohl et al., 2012).

#### 6.2 Cell culture

Cultured media was removed from each well and cells were fixed with 4% PFA for 30 min at RT. Wells were rinsed with PBS for 10 min and incubated in blocking solution (5% goat serum in 0.05% PBT) for 2 h at RT. Primary antibodies mentioned above, diluted in blocking solution, were added to each well for ON incubation at 4°C. Wells were next washed three times with PBS and incubated for 2 h at RT with appropriate Alexa-Fluor secondary antibodies, as described above. Wells were then washed and incubated with DAPI (1:400) for 15 min, washed again and kept in PBS until taken for optical analysis (Peretz et al., 2018).

### 7. Transcriptomics

#### 7.1 Library preparation and sequencing

mRNA was extracted from 6 biological replicates of FACS-isolated hindbrain cells, as described above, using Single Cell RNA purification kit (Norgen Biotek Corp., Canada), according to the manufacturer protocol. Each RNA sample had a RIN>6.9. Libraries were prepared by the Center for Genomic Technologies at the Hebrew University of Jerusalem, using the KAPA Stranded mRNA-Seq Kit (KR0960, Roche, USA) and Illumina platforms sample preparation protocol (v3.15) according to the manufacturer’s protocol. RNA sequencing was conducted by Illumina NextSeq 2000 machine using NextSeq 2000 P2, 100 cycles kit (Illumina, USA). The output was ∼ 25 million single end- 120 bp reads per sample.

#### 7.2 Bioinformatic analysis

Reads in fastq format were created with bcl2fastq v2.20.0.422, inspected for quality issues with FastQC, v0.11.8, and quality-trimmed with cutadapt, v3.4, for removal of adapters, polyA and low-quality sequences, as previously described (Alfi et al., 2021). Based on these quality analyses, reads from 4 biological replicates of the CSPG^+^ group and 6 biological replicates of the CSPG^-^ group were further analyzed by alignment to the chicken transcriptome and genome with TopHat, using genome version GRCg6a with annotations from Ensembl release 99. Quantification was done with htseq-count, v0.13.5. Differential gene expression analysis was performed using the R package DESeq2, v1.30.0 (Love et al., 2014). Genes with a sum of raw counts less than 10 over all samples were filtered out, then normalization and differential expression were calculated. Comparing CSPG+ samples to CSPG-samples was tested with default parameters using a significance threshold of padj<0.05. Whole differential expression data were subjected to gene set enrichment analysis using GSEA (Subramanian et al., 2005), (cutoff independent) in order to determine whether a priori defined sets of genes show statistically significant concordant differences between the two biological states. We used the hallmark and Gene Ontology Biological Process (GO) gene sets collections, all taken from the molecular signatures database MSigDB (Subramanian et al., 2005). A plot of protein-protein interaction network was generated, showing the interaction network between functionally important genes in the CSPG+ vs CSPG-groups. The interaction information was obtained from IPA (Ingenuity Pathway Analysis, QIAGEN Inc., https://www.qiagenbioinformatics.com/products/ingenuity-pathway-analysis) and the STRING database (Szklarczyk et al., 2023). The figure was generated using Cytoscape (Shannon et al., 2003). RNA-seq data reported in this paper was deposited in the GEO database and was given the accession number GSE230804.

### 8. Real-Time PCR

mRNA was extracted from whole hindbrains or 5-days old CSPG+ cultures, treated as described in section 2.4, using Single Cell RNA Purification Kit (#51800; Norgen Biotek Corp., Canada) according to the kit protocol, along with Norgen’s RNase-Free DNase I Kit (Norgen Biotek Corp., Canada) to degrade remaining DNA. cDNA was prepared using a High-Capacity cDNA Reverse Transcription kit (Thermo Fisher Scientific, USA), according to the manufacturer’s instructions. Real time (RT)-PCR was performed using Fast SYBR Green PCR Master Mix reagent (Thermo Fisher Scientific, USA) with the following reverse and forward primers: GAPDH-Fwd AGATGCAGGTGCTGAGTATG, Rev CTGAGGGAGCTGAGATGATAA; Sox2- Fwd TTAAGTGAAGGCGTGCTGC, Rev CCTCCTATCACTGCACCTTC; TUBB3- Fwd GACCGCATCATGAACACTTTC, Rev CGTGTTCTCCACCAGTTGAT; MAP2- Fwd CCTCCTAAATCTCCAGCAACTC, Rev CCCACCTTTAGGCTGGTATTT. 2 µl of cDNA were mixed with 10 µl SYBR mix, 7 µl ddH2O, and 1 µl of the selected primers. Real-time PCR amplification was performed using the following program: 95°C for 5 min, 40 cycles of 95°C for 15 seconds, then 60°C for 45 seconds. Results were normalized to GAPDH and analyzed using StepOne Software v2.2.2 (Applied Biosystems, USA) using ΔΔCT.

### 9. Imaging

#### 9.1 Scanning electron microscopy

Chick and mouse embryos (st. 18 and E10.5, respectively), were harvested and fixed for 1 h at RT with 2% PFA and Glutaraldehyde in 0.1M phosphate buffer pH7 and 1% sucrose (all from Sigma-Aldrich, USA). Next, the hindbrains were removed and placed on a cover slip, previously covered with poly-l-lysine (Sigma-Aldrich, USA). Samples were dehydrated in increasing ethanol concentrations (20%, 50%, 70%, 90%, 95%), then washed 4 times in 100% ethanol. The hindbrain samples were then moved to a Critical Point Dryer (Quorum K850, UK) and were gold-coated in a gold sputter coating unit (Quorum Technologies, UK). Samples were observed by low-vacuum scanning electron microscopy (SEM; JSM 5410 LV, Jeol Ltd, USA). For correlative SEM-confocal analysis, hindbrains were taken to immunofluorescence staining as mentioned above, prior to the dehydration step. Hindbrains were first imaged using a confocal microscope (Zeiss LSM-510, with Argon-Ion and 2 He-Ne Lasers), then resumed to proceed with the SEM preparation.

#### 9.2 Light and confocal microscopy

Flat-mounted hindbrains and cell cultures were imaged under an Axio Imager M1 microscope with AxioCam Mrm camera (Zeiss, Germany) or a CTR 4000 confocal microscope with DFC300FXR2 camera (Leica, Germany). Z-stack images were generated using Leica Microsystems software. For cell cultures and time-lapse analysis, an IncuCyte S3 with CMOS camera (Sartorius, Germany) was used, as previously described.

### 10. Data analysis and statistics

Fluorescence quantifications were performed using ImageJ, by subtracting the background reading out of the relative fluorescent area (Corrected Total Cell Fluorescence). For quantification of fluorescence in HBs vs. Rhs, each data point represents an average value of 4 HBs/Rhs of each embryo (n=6 for mouse, 7 for chick). For quantification of fluorescence in ChABC treated hindbrains, each bar represents an average fluorescence of area within Rhs 3-5 in treated and control embryos (n=13-20 for inhibitor injection, n=20-28 for plasmid electroporation). Average cell size in HBs and Rhs was obtained using ImageJ, each data point represents the measured area of a single cell randomly selected in different regions of the Rhs\HBs of hindbrains of two individual chick and mouse embryos. Analysis of CSPG-Sox2 proximity was done using Microscopy Image Analysis Software 9.0.2 (IMARIS; Oxford Instruments, USA). Confocal files were uploaded as 3D stacks into the software, then subjected to a 3-step workflow. First, Sox2+ cells were marked with the ‘surface’ module, defined by the estimated size of a hindbrain cell. Next, CSPG spots were segmented with the ‘spots’ module. Finally, the CSPG+ spots were linked with the surface (Sox2+ cells) with the 'spots-to-surface coloc.' script, while the inclusion criteria were set as distance between spot and surface ≤ 5um. The recorded protocol was applied for the rest of the dataset (n=7). The CSPG ‘spots’ were also used to generate a scatter plot, as each spot was assigned to its Z dimension. Cell culture analysis for neurite-length, eccentricity and floating and adherent spheres, were performed using the IncuCyte S3 live imaging system software (Sartorius, Germany). Each data point represents an average calculated in one well at day 5 of incubation. Monolayer quantification was performed using Ilastik software integrate with ImageJ analysis. Statistics were performed by unpaired t-test or one-way ANOVA followed by a Tukey post-hoc test using Graphpad Prism 8 software. P<0.05 was considered significant, data is displayed as mean±s.d.

## ACKNOWLEDGMENTS

We thank Einat Zelinger and Daniel Waiger from the Center for Scientific Imaging, Core Facility, Faculty of Agriculture, Food and Environmental Sciences, The Hebrew University of Jerusalem, for their help in the SEM and IMARIS analyses. We thank Adi Turjeman from The Center for Genomic Technologies, The Hebrew University of Jerusalem, for assisting with the RNA-sequencing. We thank Efrat Hagai from the Life Sciences Core Facilities, Weizmann Institute of Science, for assisting with the FACS procedure.

## AUTHORS’ CONTRIBUTIONS

DSD conceived the project, CH, YN, DL and AK performed the experiments. CH, YN, DL, AK and DSD analyzed the data and helped in preparing the figures; YN and HB performed the transcriptome analyses and prepared the figures, accordingly, EM provided materials.

## AVAILABILITY OF DATA AND MATERIALS

Raw RNA-seq data is available at the GEO database, accession number GSE230804. RNA-seq analyses and materials will be provided upon request.

## DECLARATIONS

Ethics approval and consent to participate

## CONSENT FOR PUBLICATION

Not applicable.

